# Multidimensional variable dose analysis (mVDA) is a novel method for high-throughput mapping of genetic interactions

**DOI:** 10.1101/2024.12.01.626212

**Authors:** S. Sengupta, B.E. Housden

**Affiliations:** Living Systems Institute, University of Exeter, Exeter, UK; Department of Clinical and Biomedical Sciences, University of Exeter, Exeter, UK

## Abstract

The mapping of genetic interactions is a powerful tool to determine gene functions, assemble the structures of biological pathways and to identify therapeutic targets for disease. However, while there have been significant advances in the screening techniques used to identify genetic interactions over the past decade, methods that are sufficiently scalable to test genetic interactions on a genome level are still far from our current capabilities. Here, we describe an approach to genetic interaction screening in *Drosophila* cells that overcomes the scaling issues associated with most other methods. This method, called multidimensional Variable Dose Analysis (mVDA), allows multiple, random genes to be inhibited within each cell of a mixed population and the relative phenotypes caused by each gene or pair of genes to be deconvoluted. This means that reagent library size and cell population size do not scale exponentially with the number of genes to be tested, unlike previous methods. This method therefore has the potential to allow genome wide mapping of genetic interactions in *Drosophila* cells for the first time.

## Introduction

The mapping of genetic interactions in which the function of one gene influences the function of another has been a goal of the functional genomics field for many years. Previous studies have established that knowledge of genetic interactions can generate significant jumps in our understanding of biology. For example, genetic interactions studies have enabled the assembly of the fundamental pathways controlling key biological processes such as the cell cycle (Hartwell *et al*. 1970; Hereford and Hartwell 1974), cell polarity (Labbe *et al*. 2006) and core signalling pathways (Simon 1994; Verheyen *et al*. 1996; St Johnston 2002; Collins and Cohen 2005; Lehner *et al*. 2006; Horn *et al*. 2011; Geisbrecht *et al*. 2013). More recently, genetic interaction screens have enabled identification of therapeutic drug targets for cancers (Zhou *et al*. 2014; Housden *et al*. 2015; Housden *et al*. 2017a; Valvezan *et al*. 2017; Nicholson *et al*. 2019; Previtali *et al*. 2024; Stevens *et al*. 2024) and functional annotation of many individual genes (Tong *et al*. 2004; Housden *et al*. 2015; Costanzo *et al*. 2016; Stevens *et al*. 2024).

A major challenge in this field is to develop genetic screening methods that can be scaled sufficiently to map genetic interactions across whole genomes in complex model systems. So far, large-scale genetic interaction mapping has only been possible in relatively simple model systems such as the yeast *Saccharomyces cerevisiae* (Tong *et al*. 2004; Costanzo *et al*. 2010). In more complex models such as *Drosophila* and humans, mapping of genetic interactions has been far more limited in scale (Horn *et al*. 2011; Fischer *et al*. 2015; Han *et al*. 2017; Shen *et al*. 2017; Horlbeck *et al*. 2018). While databases such as <u>flybase</u> have accumulated many different annotations of genetic interactions, it is clear that interactions are not constant between cell types or environmental conditions (Horn *et al*. 2011; Housden *et al*. 2017b; Ryan *et al*. 2018). Therefore, it is necessary to map interactions at a large-scale under controlled conditions to maximise the insight that can be gained. This further emphasises the need for easily scalable screening methods because interactions will need to be assessed repeatedly under differing conditions in order to understand how they are altered by cell states. For example, knowledge of how the genetic interaction network differs between healthy and diseased states will likely result in many novel discoveries relevant to establishing effective therapies.

*Drosophila* cells are a well-established model for genetic screening with many hundreds of screens having been performed across a wide range of research fields (Mohr *et al*. 2014; Housden *et al*. 2017b; Hu *et al*. 2021; Viswanatha *et al*. 2024). In addition, many genes are well conserved between *Drosophila* and humans with approximately 65% of disease associated genes having *Drosophila* orthologs (Chien *et al*. 2002; Yamamoto *et al*. 2014; Ugur *et al*. 2016). Furthermore, *Drosophila* models have been established for many different human diseases illustrating the relevance of this model for understanding human biology and developing new therapies for disease (Mohr; Chien *et al*. 2002; Yamamoto *et al*. 2014; Housden *et al*. 2015; Ugur *et al*. 2016; Nicholson *et al*. 2019; Stevens *et al*. 2024).

The *Drosophila* genome has reduced complexity compared to the human genome. For example, there are fewer genes and fewer redundancies between gene functions. While this may mean that not all aspects of human biology can be studied in *Drosophila*, it makes the comprehensive mapping of genetic interactions more feasible. Genetic interaction screen size scales exponentially with the number of genes included. Combined with the well-established and robust genetic tools available in *Drosophila*, the simpler genome therefore makes this an excellent system in which to develop new approaches for genetic interaction mapping that can later be converted for use in human cells.

Here, we describe a new approach to viability measurement that relieves the limits on scaling by allowing multiple conditions to be assessed in the same cells. We show that genetic interactions can be accurately identified from these assays and that the method has high potential for scaling to allow whole genome mapping of genetic interactions in *Drosophila* cells for the first time.

## Results

### Multidimensional Variable Dose Analysis (mVDA) is a novel assay with the ability to deconvolute phenotypes from mixed RNAi reagents

As described above, a key limitation of existing genetic interaction screening methods is an inability to efficiently scale reagent libraries and cell populations to allow genome scale interrogation of gene pairs. This lack of scalability stems from the fact that for all but one existing pooled screening method (Dixit *et al*. 2019), different genes or pairs of genes must be inhibited in each individual cell within a mixed population such that the loss of cells can be attributed to the reagent that they contained. This enables screens to be performed in which sequencing of the CRISPR or RNAi reagents enables inference of viability phenotypes due to the relative loss of specific reagents or reagent pairs from the cell population due to the death of the cells that contained them.

To enable improved scaling of genetic interaction screens, we investigated whether it would be possible to deliver multiple RNAi reagents to each cell in such a way that their relative contribution to cell death could be determined from the cells remaining.

We previously developed an assay called Variable Dose Analysis (VDA) in *Drosophila* cells, in which shRNA-encoding plasmids are transfected into a population of cells with a range of doses delivered to each individual cell (Housden *et al*. 2017a; Sierzputowska *et al*. 2018; Wang and Housden 2024). Cell viability can then be determined by analysing the distribution of reagent doses within the surviving population. Reagents that cause a loss of cell viability produce a shift in this distribution towards lower reagent doses due to more efficient induction of cell death by high RNAi dose (**Figure 1A**). This method was initially developed to enable analysis of the phenotypes of essential genes at sub-lethal knockdown efficiency (Housden *et al*. 2017a). An additional advantage of the method is that the signal to noise ratio is improved by approximately 12-fold compared to previous dsRNA-based methods (Wang and Housden 2024).

**Figure 1:**
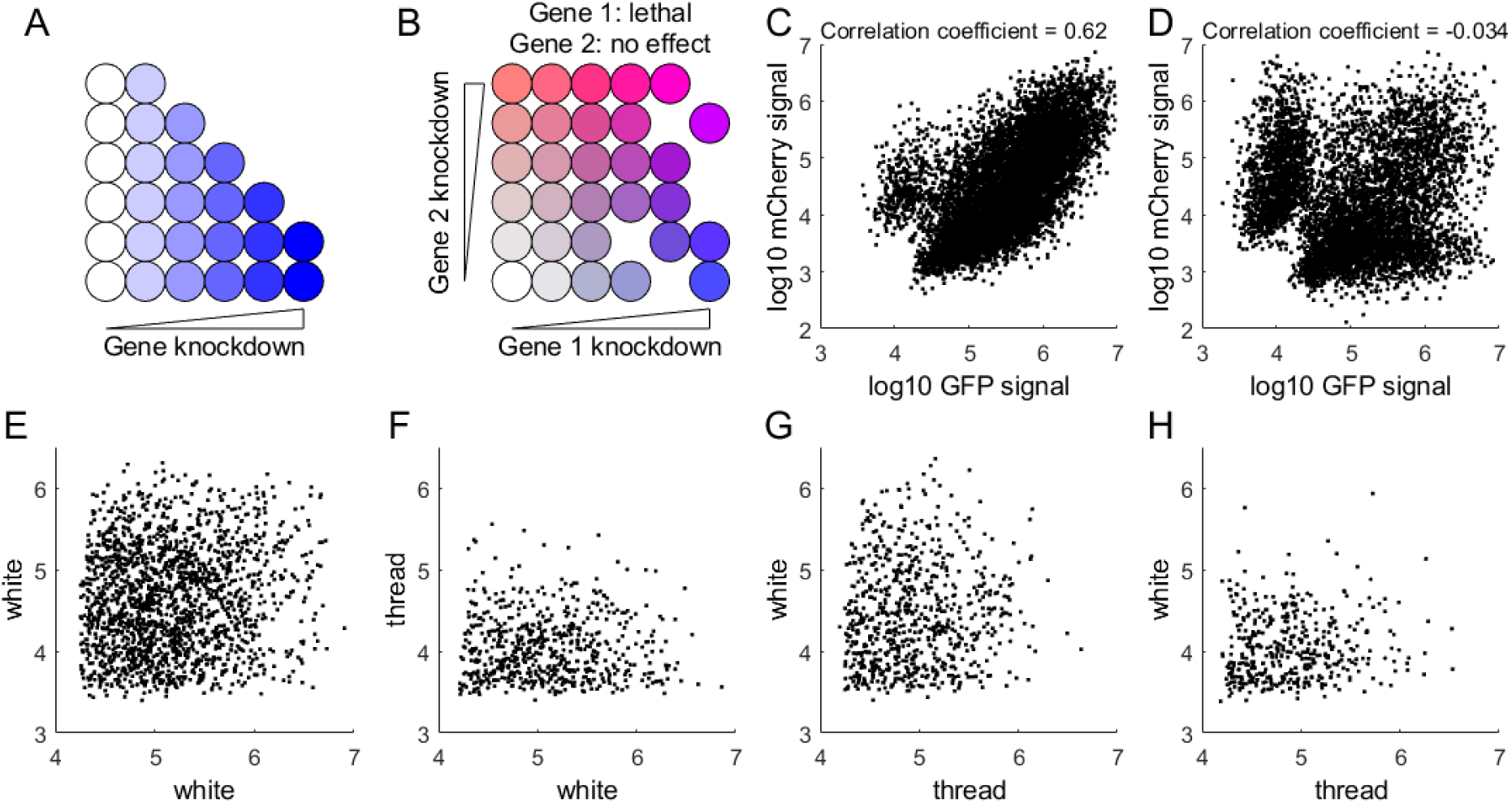
mVDA assays can be multiplexed while maintaining independent signals. **A)** Diagram illustrating how viability phenotypes are determined based on analysis of the distribution of cell numbers in relation to knockdown efficiency of an essential target gene in VDA assays. Cells with greater knockdown of the target gene are preferentially lost resulting in a change in dose distribution. **B)** Diagram illustrating how viability phenotypes can be determined for two different target genes using a multiplexed VDA (mVDA) assay. Viability effects from gene 1 have no effect on the distribution of cells in the dimension associated with gene 2. **C-D)** Scatter plots showing the correlated or decoupled delivery of transfected plasmids. Each dot represents the GFP and mCherry fluorescent signal from a single cell in a transfected population. Data are a representative example from two biological replicates and correlation coefficients were calculated from the combined data from both replicate experiments. **E-H)** Experimental data from two-dimensional mVDA assays with each dimension reported using GFP (horizontal axis) or mCherry (vertical axis). Data show the distribution of fluorescent signals (log10 of fluorescent signal) for each dimension and illustrate the independence of changes in distribution between the two dimensions. ‘white’ and ‘thread’ indicate the target of the shRNA reagent used in the relevant dimension.

We have used VDA for the discovery of candidate drug targets for both Tuberous Sclerosis Complex and Neurofibromatosis type 1 (Housden *et al*. 2017a; Stevens *et al*. 2024), validating the ability of the method to assess changes in cell viability due to inhibition of target genes.

We hypothesised that it would be possible to transfect multiple shRNA plasmids to the same cells and separate the resulting viability phenotypes so long as the distribution of doses of each plasmid were independent. In this case, a change in distribution for one reagent would not affect the distribution of the other, which is represented in an independent dimension within the cell population (**Figure 1B**). We termed this method multidimensional VDA (mVDA).

To test the underlying requirement of the mVDA method, that shRNA reagent can be delivered independently to a cell population, we transfected cells with plasmids encoding either a GFP or mCherry reporter. We used either a standard co-transfection method in which the plasmids are mixed and then transfected or using an alternative method in which transfection complexes are formed independently for each plasmid and then delivered separately to the cells. The relationship between the dose of each plasmid delivered to each cell was then assessed using cytometry to measure the distribution of GFP and mCherry fluorescence. We found that while the standard co-transfection method produced a clear relationship between the dose of plasmids delivered to each cell (**Figure 1C**), the alternate method resulted in independent doses with a correlation coefficient close to zero (**Figure 1D**). This result indicates that by delivering transfection complexes separately it is likely possible to generate the independent distributions of shRNA reagents required for the mVDA method.

Next, we tested whether similar independent distributions could be obtained in the presence of shRNA-encoding plasmids targeting control genes with known effects in cell viability. Transfections were performed in which shRNA targeting the *white* gene, which is often used as a negative control in RNAi-based experiments, was mixed with reporter plasmids expressing either GFP or mCherry. These plasmid mixtures were then transfected independently into a population of cells and the fluorescent signals assessed. As observed with the fluorescent reporters alone, the doses of fluorescent reporter plasmids received by each cell were independent (**Figure 1E**).

We further considered cases in which one or both of the shRNAs targeted a gene known to cause reduced viability when inhibited (*thread*, an apoptosis inhibitor known to induce efficient cell death when inhibited). In each case, the distribution of the fluorophore associated with the shRNA targeting *thread* was altered relative to the control distribution but the fluorophore associated with the shRNA targeting *white* was unaffected (**Figure 1F-G**). Similarly, when both GFP and mCherry were associated with the *thread* shRNA, both distributions were affected (**Figure 1H**). These results indicate that the viability effects associated with particular shRNA reagents can be resolved from a mixed population of cells in which multiple shRNAs are present.

### The mVDA method can be scaled to many mixed RNAi reagents to enable screening applications

A key feature of the mVDA method is that it has the potential to scale to many dimensions. While our initial experiments demonstrate that viability effects can be measured from populations of cells containing two dimensions, we sought to assess whether higher numbers of dimensions would be possible.

To test this, we set up experiments in which the *white* and/or *thread* shRNAs were transfected independently into cells with between two and twenty dimensions. In each case, we considered four different conditions: 1) all dimensions received the *white* shRNA (**Figure 2A, black bars**), 2) one dimension received *thread* shRNA and all other dimensions received *white* shRNA (**Figure 2A, yellow bars**), 3) one dimension received *white* shRNA and all other dimensions received *thread* shRNA (**Figure 2A, blue bars**) or 4) all dimensions received *thread* shRNA (**Figure 2A, white bars**). In all of these experiments, only the first dimension was marked with GFP and the other channels were unmarked.

**Figure 2:**
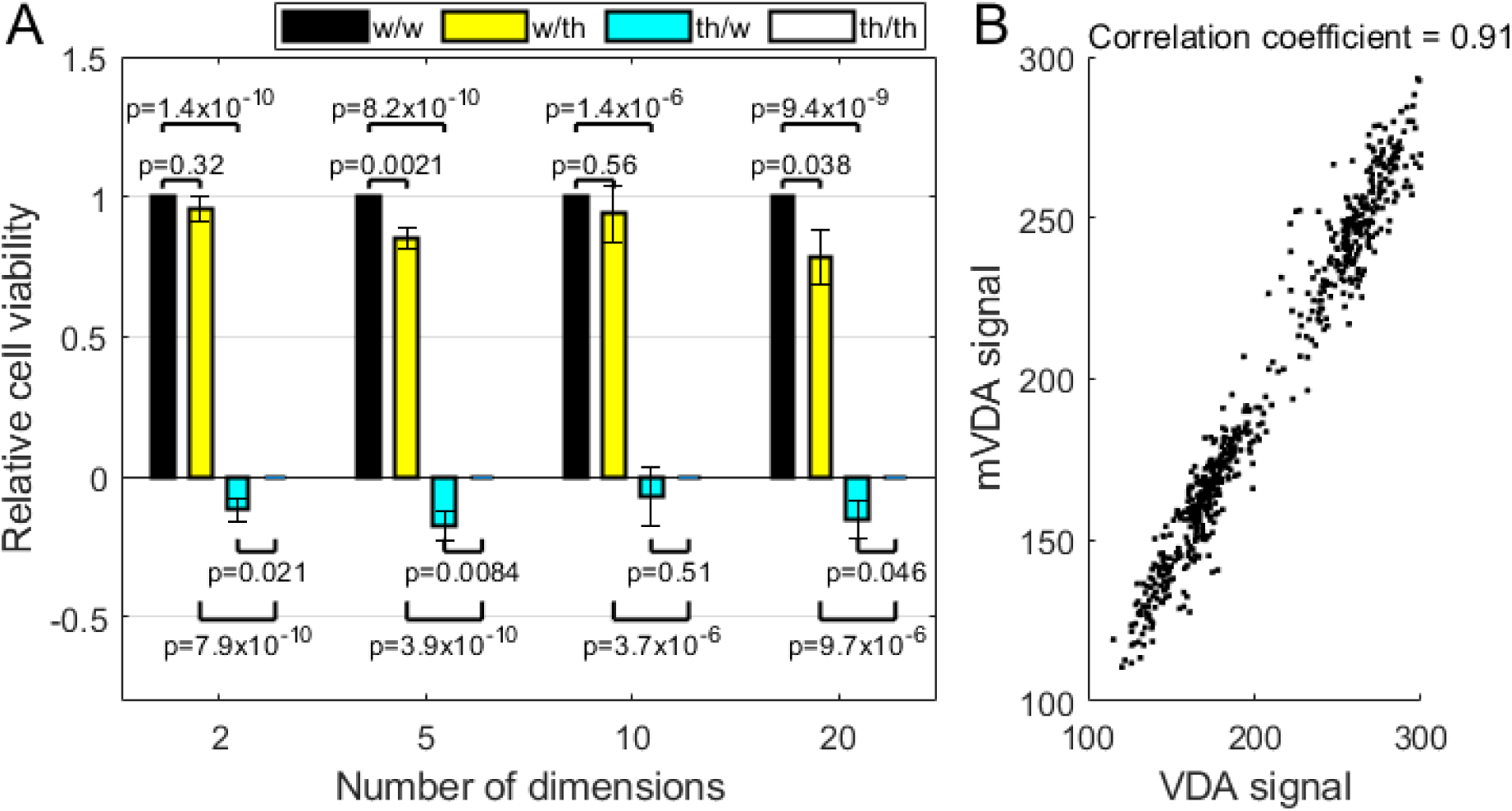
mVDA accurately reflects viability phenotypes. **A)** Bar graph showing the viability effect caused by a single shRNA reagent multiplexed with a total of 2, 5, 10 or 20 dimensions per experiment, as indicated. Viability is illustrated as VDA signal for the relevant dimension normalised to negative and positive control conditions such that the negative control =1 and the positive control =0. For the negative control (black bars), all dimensions are associated with the ‘white’ shRNA and for the positive control (white bars), all dimensions are associated with the ‘thread’ shRNA. For each condition, the legend indicates the shRNA associated with the dimension measured (listed first) and the shRNA associated with all other dimensions (listed second). Each bar represents the average of at least five biological replicates and error bars indicate standard error of the mean. P-values were calculated using unpaired, two-tailed t-tests. **B)** Scatter plot comparing viability phenotypes measured using VDA or mVDA for approximately 450 shRNA reagents.

The distributions of GFP within the various cell populations were measured using cytometry. Results were normalised to the *white* only (condition 1) and *thread* only (condition 4) data. We found that in all cases, the marked channel corresponded closely with the expected result from the shRNA included in that dimension. All samples in which *white* shRNA was measured produced viability signals close to the negative control and all samples in which *thread* was marked produced signals close to the positive control (**Figure 2A**). We note that some samples produced signals that were significantly different from the relevant negative control (e.g. 20-dimensions, condition 1 compared to condition 2) although these differences were much weaker than those between positive and negative control reagents (e.g. 20-dimensions condition 2 and condition 4). These results suggest that the method is highly scalable and that even very strong viability phenotypes (e.g. those produced by 19 dimensions expressing *thread* shRNA) have little effect on phenotypes measured from independent dimensions.

To assess the ability of mVDA to measure viability phenotypes more broadly, we transfected approximately 450 shRNAs organised randomly into pairs and each marked by GFP or mCherry and measured viability phenotypes through the fluorophore distributions as previously described (Housden *et al*. 2017a; Sierzputowska *et al*. 2018; Wang and Housden 2024). In parallel, we transfected each shRNA individually such that ground-truth viability effects could be determined. Comparison between viability effects showed a correlation coefficient of 0.91, indicating that the multiplexed method accurately measures cell viability (**Figure 2B**).

### mVDA accurately identifies genetic interactions

So far, we have shown that mVDA assays can distinguish between signals produced by single shRNA reagents in a mixed population transfected with multiple shRNAs. However, the greatest potential advantage of mVDA comes from the possibility of measuring interactions between genes from the same mixed population.

To test this, it was first necessary to identify control genetic interactions. Genetic interactions are highly variable between cell types. In addition, previous methods for mapping genetic interactions in *Drosophila* cells used a different approach to assessing viability phenotypes (e.g. dsRNA). Therefore, to identify robust interactions, we first tested nine interactions that have been demonstrated in S2R+ cells from previous experiments using various methods. We then tested these interactions using VDA assays in which pairs of shRNAs were co-transfected with correlated distributions. Of the nine interactions, four produced significantly different results when combined compared to the expected result determined based on the effects of inhibiting each gene alone (**Figure 3A**). These four gene pairs were considered to be ‘true’ genetic interactions for the purpose of testing the mVDA method.

**Figure 3:**
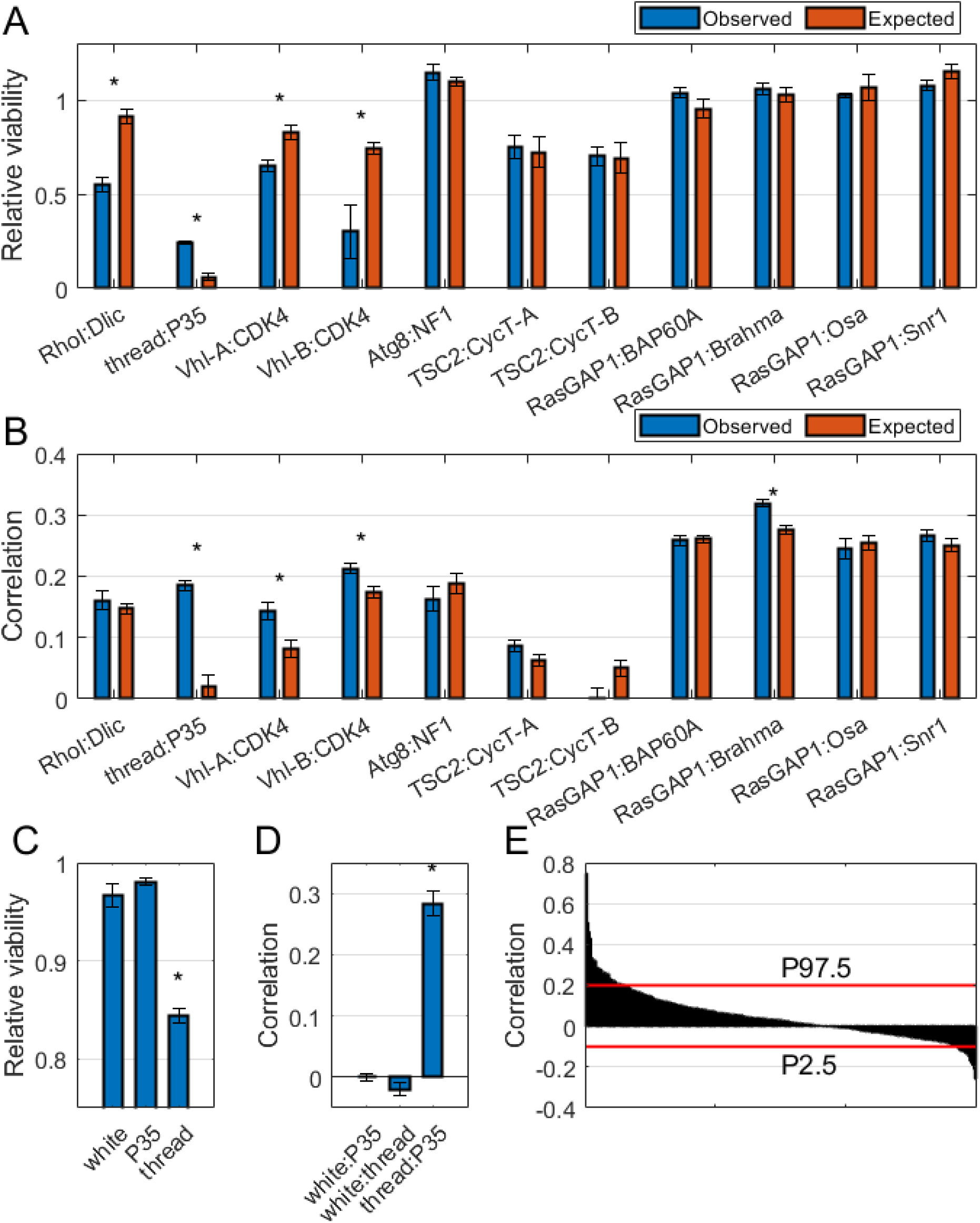
mVDA accurately detects genetic interactions. **A)** Bar chart showing the observed (blue) and expected (red) viability of S2R+ cells transfected with the indicated pairs of shRNA reagents and analysed using VDA assays. Each bar represents the average of at least 12 replicates and error bars indicate standard error of the mean. Asterisks indicate p<0.05 based on Mann-Whitney U tests. **B)** Bar chart showing the observed (blue) and expected (red) viability of S2R+ cells transfected with the indicated pairs of shRNA reagents, each delivered to an independent dimension within the cell population. Each bar represents the average of at least 12 replicates and error bars indicate standard error of the mean. Asterisks indicate p<0.05 based on Mann-Whitney U tests. **C)** Viability effects determined from a three-dimensional mVDA experiment. Bars represent the average of 24 replicates and error bars indicate standard error of the mean. The asterisk indicates p<0.05 based on Mann-Whitney U tests. **D)** Correlation coefficients from the indicated pairs of reagents in a three-dimensional mVDA experiment. Bars represent the average of 24 replicates and error bars indicate standard error of the mean. The asterisk indicates p<0.05 based on Mann-Whitney U tests. **E)** Linear correlation coefficients from pairs of shRNA reagents assessed with two-dimensional mVDA assays. These data are from the same experiment illustrated in Figure 2B. Red lines indicate the cutoffs for the top and bottom 2.5% of the distribution of correlations.

Next, we tested the same nine gene pairs using mVDA assays and again calculated expected and observed relationships between fluorophore distributions (see materials and methods). Of the nine pairs, four showed evidence of genetic interactions and three of these were common with the VDA assays (**Figure 3B**). This suggests that relationships between dimensions in mVDA assays are an accurate representation of genetic interactions.

One limitation of testing genetic interactions in two-dimensional mVDA assays is that it is also necessary to assess each shRNA alone to determine the expected relationship between dimensions for comparison with the observed relationship. This limits the scalability of the approach as each shRNA needs to be tested both alone and in combination with other candidate shRNAs. To overcome this, we tested whether three-dimensional mVDA assays would allow simultaneous analysis of negative control interactions in the same cell population as the candidate gene pair, thereby increasing the efficiency of the method.

shRNAs targeting *white* and *thread* as well as a plasmid overexpressing p35 protein were transfected into a single population of cells. *Thread* causes a reduction in cell viability that can be blocked via a genetic interaction with *p35*. These reagents were each marked by GFP, mCherry or a plasmid expressing a cell surface marker that could be detected with a fluorescent antibody. Fluorescence distributions were then assessed using cytometry. First, we assessed the viability effects caused by each gene individually and found that results were as expected with *white* and *p35* causing no effect and *thread* reducing cell viability (**Figure 3C**).

Next, we compared the linear relationships between each pair of fluorophore distributions. Both *white* and *p35* as well as *white* and *thread* pairs produced independent distributions indicated by a correlation close to zero. However, the *thread* and *p35* combination showed a clear relationship between distributions, with a positive correlation, indicating a genetic interaction (**Figure 3D**).

To build on this and assess the ability of mVDA to identify genetic interactions more broadly, we reanalysed the data from the 450 gene pairs assessed previously (**Figure 2B**) to identify genetic interactions between each pair of genes. We then ranked all of the gene pairs based on the correlation coefficient between the paired distributions (**Figure 3E**). To validate the identified interactions, we assessed all pairs of genes that are conserved in yeast to determine whether genetic interactions were also identified in a large scale genetic interaction map previously generated in this system (Costanzo *et al*. 2010). This analysis showed that the top and bottom 2.5% of ranked pairs were highly enriched for genetic interactions also found in the yeast screens. Specifically, from the dataset, 67/207 gene pairs were conserved in yeast and present in the yeast dataset. Of these, 12, were identified in the top and bottom 2.5% of the mVDA results and 9 of those (75%) also had genetic interactions in the yeast database. Conversely, only 22/55 (40%) of the remaining conserved gene pairs had genetic interactions in the yeast database. These results indicate that mVDA accurately identifies genetic interactions.

### The mVDA method is highly scalable and has potential to allow genome scale mapping of genetic interactions

The number of genetic interactions that can be simultaneously tested using mVDA assays increases exponentially with the number of dimensions. Therefore, the power of the mVDA screening method is dependent on the number of dimensions that can be assessed in a single population of cells. To test the scaling of dimensionality in mVDA assays, we performed an experiment in which 736 different transfections were performed independently into a single population of cells. This number of dimensions potentially allows approximately 270,000 genetic interactions to be measured simultaneously, rivalling the largest genetic interaction screens ever performed in metazoan models.

The majority of the 736 dimensions included the *white* shRNA and no fluorescent reporter. Three dimensions contained *white* shRNA, *thread* shRNA and *p35*, each marked by a different fluorescent reporter. Cytometry analysis was then performed and relationships between distributions assessed. Considerable variation was observed in the correlation between distributions, preventing direct statistical analysis of results between replicates. However, when the genetic interactions were ranked, the expected interaction between *thread* and *p35* ranked top in all replicate experiments. This indicates that while scaling of the number of dimensions does increase quantitative variation, genetic interactions can be reproducibly identified through ranked data (**Figure 4A**).

**Figure 4:**
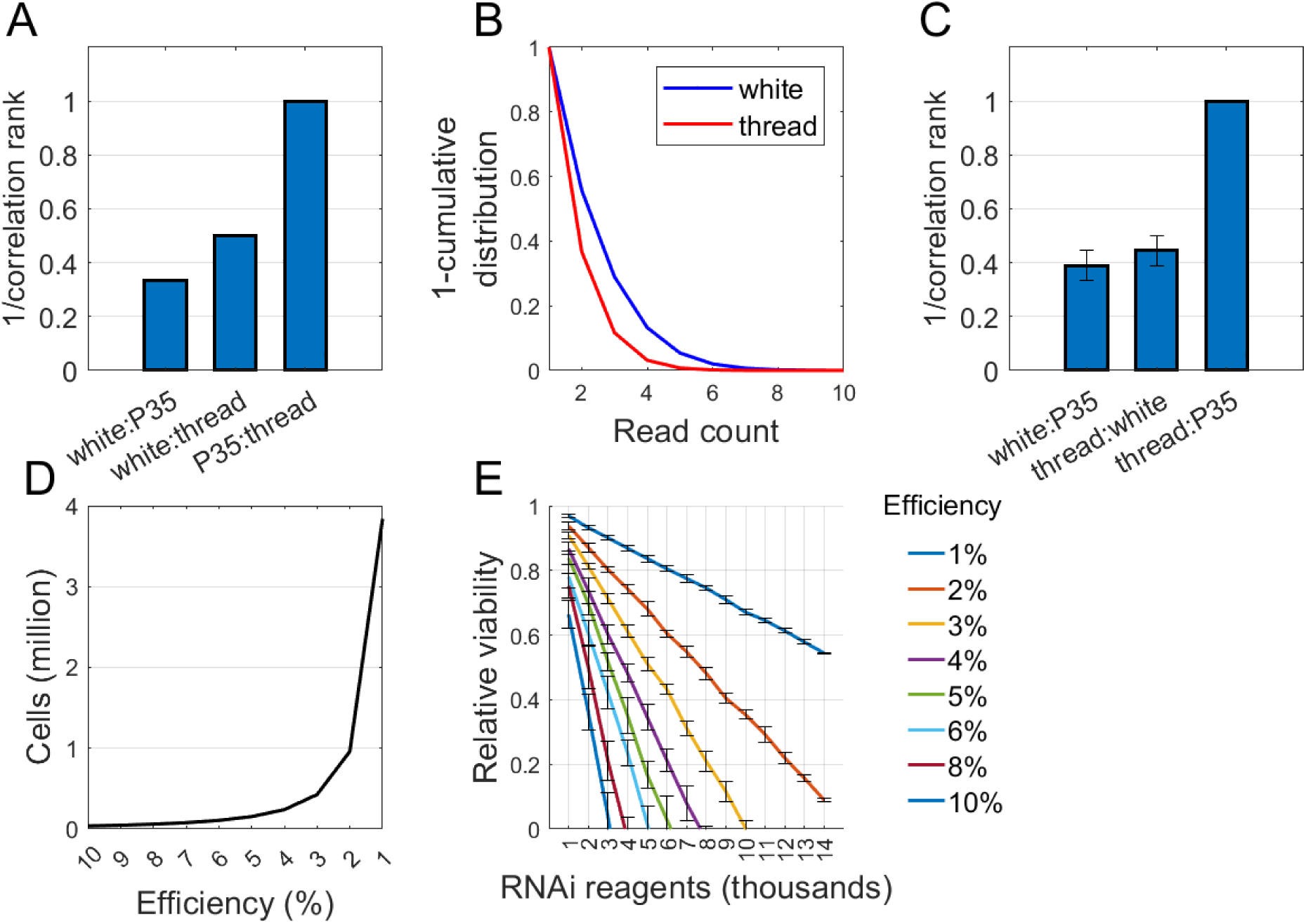
The mVDA method is highly scalable. **A)** Bars indicate the average 1/ranked correlation coefficient from three replicate 736-dimension mVDA experiments. Error bars indicate standard error of the mean and equal zero due to consistent ranking between replicates. **B)** Lines represent 1-cumulative distributions of read counts per cell associated with either white (blue line) or thread (red line) shRNA. The graph represents example data from three replicate experiments. **C)** Bars indicate the average 1/ranked correlation coefficient from three replicate 3-dimension mVDA experiments assessed using single cell sequencing. Error bars indicate standard error of the mean**. D)** Prediction of the relationship between the efficiency of transfection (proportion of cells receiving each shRNA reagent) and the number of cells that must be sequenced to identify genetic interactions. **E)** Prediction of the relationship between cell population viability and the number of reagents screened (dimensions) for different transfection efficiencies.

These results suggest that the mVDA method is a powerful tool to identify genetic interactions and is highly scalable due to the ability to assess multiple shRNAs in a single population of cells. However, the number of dimensions that can be assessed in a single experiment is limited by the number of available genetically encoded fluorescent reporters and the ability of cytometry instruments to differentiate fluorescent signals. This readout therefore limits the number of dimensions to approximately 50 as a maximum with existing cytometry technology. To address this issue, we investigated whether the cytometry readout could be replaced with a single cell sequencing method to assess the relative levels of each shRNA encoding plasmid between cells in a transfected population. This has the added advantage of not requiring co-delivery of the shRNA and fluorescent reporter plasmid, likely reducing noise.

As an initial test, we performed single-dimension VDA assays using the *white* and *thread* shRNA plasmids. We then performed single cell sequencing using a split-pool barcoding approach that can potentially be used to label an unlimited number of cells (see materials and methods). This scaling ability will be necessary when the number of dimensions are increased to perform larger screens. Sequencing results were analysed in a similar way to the previous cytometry results to produce distributions of shRNAs across the cell population. A clear difference between the distributions of *white* and *thread* shRNAs was identified, indicating that single cell sequencing can reliably identify viability effects represented by changes in the distribution of shRNA encoding plasmids, similar to the cytometry-based readouts (**Figure 4B**).

To determine whether the single-cell sequencing readout could detect genetic interactions, we performed a three-dimensional experiments in which *white*, *thread* and *p35* were included in independent dimensions. Similar to the previous cytometry-based experiments, ranking of the correlations between distributions resulted in reproducible detection of the genetic interaction between *thread* and *p35*, with this gene pair consistently ranking top in all replicates (**Figure 4C**).

To assess the likely limits of scalability of the mVDA method, we produced a computational model of the method, taking into account several experimentally determined the parameters. The distribution of shRNA delivery within a population of cells was determined based on fluorescence distributions from the cytometry experiments and the effects of RNAi treatments on cell viability were based on results from previous whole genome RNAi screens for cell viability performed using S2R+ cells (Stevens *et al*. 2024). Finally, the number of cells required to accurately detect a genetic interaction was based on the experiments described above.

We varied two parameters within this model to assess both the number of cells that would need to be sequenced to analyse all pairwise combinations of genes and the number of dimensions that could be included without the burden of viability phenotypes causing widespread cell death.

The results from this model indicate that the approach is highly scalable. The cumulative viability phenotypes eventually limit the number of shRNAs that can be delivered to each individual cell (**Figure 4G**). However, this can be accounted for simply by limiting the proportion of cells in the population that receive each reagent, thereby reducing the shRNA burden in each cell. The disadvantage of limiting the delivery in this way is that more cells must be sequenced in order to gain sufficient distribution data for each pairwise combination of genes (**Figure 4F-G**). Based on the model, we predict that up to 9,000 dimensions are expected to be possible with each shRNA delivered to 2% of the cell population, to prevent population viability loss. The model also predicts that this condition would require sequencing of approximately one million cells, which is well within the ability of current single cell sequencing methods. In particular, the iterative barcoding method that we previously employed (**Figure 4B**) enables the labelling of a very large number of cells simply by performing additional rounds of iterative barcoding. Therefore, the number of cells to be sequenced is unlikely to prevent large-scale screens although this would increase the cost of such screens.

Overall, this analysis suggests that the mVDA method is highly scalable and has a high chance of enabling analysis of 9,000 shRNA reagents, representing over 400 million genetic interactions in a single experiment. This is around 1,000-fold higher in scale than the largest genetic interaction screens performed to date in either *Drosophila* or human cells.

## Discussion

In this study, we have developed a novel approach to mapping genetic interactions, called mVDA. The key advantage of this method is that shRNA reagents can be delivered to a population of cells with no need to maintain separate sub-populations for each reagent or pair of reagents to be tested. This overcomes a major limitation of most previous methods. In addition, previous methods required that a new reagent be generated for each combination of genes to be inhibited to ensure co-delivery to a distinct subset of cells. This means that the reagent library and transfection scale needs to be increased exponentially as the number of genes tested is increased. Similarly, as each reagent or pair of reagents requires an independent subset of cells within the population, the number of cells also increases exponentially with screen size. By contrast, the mVDA method allows delivery of multiple shRNA reagents to each cell in the population with random combinations. The relative contribution to viability phenotypes is then determined by analysing the interactions between distributions of reagents within surviving cells. Therefore, it is not necessary to increase either reagent library size or cell population exponentially with increasing screen scale. Indeed, our modelling suggests that the scaling of the number of cells that must be transfected and sequenced is far below exponential in nature.

To allow large-scale mapping of genetic interactions using mVDA, the next steps will be to first generate genome-wide libraries of shRNA encoding plasmids compatible with this method. While large shRNA reagent libraries already exist (Zirin *et al*. 2020), these currently require the use of Gal4 to drive expression. However, these reagents can potentially be subcloned relatively easily to generate compatible libraries in the future.

Despite the success of the mVDA method in identifying genetic interactions in this study, there remain multiple aspects of the method that could be optimised. For example, the optimal number of cells and sequencing reads per cell have not yet been determined. In addition, it has previously been shown that genetic interaction strength does not always increase as the knockdown level of the target genes in increased (Horn *et al*. 2011). The current analysis method relies on linear correlation, which will be more sensitive to interactions that increase linearly with knockdown strength. Therefore, more complex analysis methods must be developed to maximise identification of different classes of genetic interaction.

It may be necessary to optimise further as the method is scaled up. Our modelling suggests that scaling to many thousands of dimensions will be possible before anticipated limits are reached in terms of accumulated viability phenotypes. Further optimisation will likely allow this to be increased even further to allow full-genome screens for the first time.

Cell viability may be affected by cell cycle phenotypes as well as cell death. We would predict that a slowing of the cell cycle may lead to accumulation of larger cells and a reduction in the dilution of the shRNA-encoding plasmids by cell division. This may cause a shift in distribution towards higher doses, the opposite of the cell death phenotype (and we have observed this anecdotally). Additionally, knockdown of a single gene may lead to both of these phenotypes at different knockdown efficiencies, creating even more complex changes to the distribution. These issues may result in the failure to identify genetic interaction however they also represent a valuable opportunity to extract more nuanced information about gene functions from the screen data. Again, this highlights the need to develop more advanced ways of assessing the reagent distributions both to enhance detection of genetic interactions as well as to characterise those interactions more precisely.

In the future, it will be valuable to apply this screening method to human cells to allow analysis of genes and interactions that are not conserved in *Drosophila*. In addition, it will likely be valuable to explore how genetic interactions vary between different human cell types, which cannot be done using the *Drosophila* system. We expect that the method will be applicable in human cells with minimal alteration. However, not all human cell lines are easily transfected. A key requirement for the mVDA method is that a broad range of plasmid doses are delivered across a cell population and that these dose distributions are independent for each reagent. Poor transfection efficiency in some human cell lines may make this challenging. Further work will be needed to overcome these potential issues.

Finally, we note that a key future possibility of this method is the ability to analyse higher order genetic interactions without the need to greatly increase screen scale. As the shRNA reagents are already delivered to the same population of cells in pairwise genetic interaction screens using the mVDA method, it will likely also be possible to extract information on interactions between three or more reagents. The key factor determining this will be the number of shRNAs delivered to each individual cell and the number of cells containing each specific mixture of reagents. Therefore, some scaling-up will be required although this is likely to be to a much lower extent that would be needed using more traditional screening approaches.

## Methods

### *Drosophila* cell culture

S2R+ cells were cultured using standard methods. For cell maintenance, cells were incubated at 25 °C in Schneider’s medium supplemented with 10% FBS and 5% penicillin/streptomycin.

### Transfections

Transfections were performed using Fugene HD transfection reagent (Promega) according to manufacturer’s recommendations. A ratio of 3:1 of transfection reagent to DNA was used for all experiments. For VDA assays, 190 ng of shRNA plasmid were mixed with 10 ng of fluorescent reporter plasmid before addition of 0.6 µL of the transfection reagent in a total volume of 10 µL per well of a 96-well plate. For mVDA experiments a similar approach was taken except that transfection mixtures were generated independently and only mixed on addition to the cell cultures.

### Flow cytometry

Flow cytometry was performed using a Beckman Coulter Cytoflex S system with a 96-well plate loader. Data were analysed using Cytexpert (Beckman Coulter) to define fluorescent cells and then further analysis was performed using custom Matlab scripts.

### Single cell sequencing

For the three-dimensional mVDA experiments, single cells were isolated using a FACSAria system to sort single GFP positive cells into 384-well plates. Cells were then labelled using PCR to amplify a unique barcode sequence of the transfected shRNA encoding plasmids, including unique cell barcode sequences on the PCR primers. Sequencing was then performed using a Novaseq platform. Data were analysed using Matlab to identify and count barcode sequences and cell label sequences. Distributions of reads from each cell were then compared using linear correlation to identify genetic interactions.

A split-pool barcoding method was also used to establish an approach to scale up the single cell sequencing (e.g. Figure 4B). The principle of this method is described in a previous publication (Rosenberg *et al*. 2018). However, in our method, iterative barcoding was performed using PCR rather than ligation. In each round, 96 different 4-bp barcode sequences were added to the cell population before re-pooling and repeating the barcoding process. A total of four rounds of barcoding were used, leading to approximately 85 million distinct cell labels.

### Validation of genetic interactions using yeast data

Comparisons with the yeast database of genetic interactions (Costanzo *et al*. 2010) were performed by first mapping orthologs of the tested genes in *Saccharomyces cerevisiae* using the DIOPT tool (Hu *et al*. 2011) using a DIOPT score cutoff of 3. Next the yeast database available at thecellmap.org was manually searched for genetic interactions for each gene pair. Reporting of a genetic interaction in this tool taken as a positive result with no quantitative cutoff applied.

### Modelling

The model is based on the following assumptions, which are informed by experimental data: 1) Transfection results in a normal distribution of shRNA delivery across a population of cells. 2) The proportion of essential genes and strength of viability phenotypes will be similar to previous genome-wide RNAi screens (Stevens *et al*. 2024). 3) Viability phenotypes will be additive within each individual cell and scale linearly with dose of the shRNA reagent. 4) 384 cells containing both of a pair of shRNA reagents are required to identify a genetic interaction. Each datapoint in Figure 4E is the average prediction of 1,000 cycles of modelling in which shRNAs are delivered to random subsets of the cell population. Error bars indicate standard deviation.

### VDA assays

Fluorescent signals from VDA assays were analysed as described previously (Sierzputowska *et al*. 2018). For genetic interaction experiments, expected viability scores were determined by summing the differences between the negative control (*white*) and each of the tested genes alone and adding this from the negative control viability.

### mVDA assays

mVDA assays were performed as described in the transfection section to generate independent distributions of shRNA-encoding plasmids within cell populations. Data were collected either using flow cytometry to assess the expression of multiple fluorophores or using single cell sequencing to quantify the relative levels of each shRNA plasmid in each cell. Genetic interactions were identified using linear correlation analysis. Expected correlation coefficients were determined by averaging the correlation coefficients between the distributions of the negative control shRNA (*white*) and each experimental shRNA alone.

